# Personalised metabolic whole-body models for newborns and infants predict growth and biomarkers of inherited metabolic diseases

**DOI:** 10.1101/2023.10.20.563364

**Authors:** Elaine Zaunseder, Ulrike Mütze, Jürgen G. Okun, Georg F. Hoffmann, Stefan Kölker, Vincent Heuveline, Ines Thiele

**Affiliations:** Engineering Mathematics and Computing Lab (EMCL), Interdisciplinary Center for Scientific Computing (IWR), Heidelberg University, Heidelberg, Germany; Data Mining and Uncertainty Quantification (DMQ), Heidelberg Institute for Theoretical Studies (HITS), Heidelberg, Germany; School of Medicine, University of Galway, Galway, Ireland; Division of Child Neurology and Metabolic Medicine, Center for Child and Adolescent Medicine, Heidelberg University Hospital, Heidelberg, Germany; Discipline of Microbiology, University of Galway, Galway, Ireland; Ryan Institute, University of Galway, Galway, Ireland; APC Microbiome Ireland, Cork, Ireland

## Abstract

Extensive whole-body models (WBMs) accounting for organ-specific dynamics have been developed to simulate adult metabolism. However, there is currently a lack of models representing infant metabolism taking into consideration its special requirements in energy balance, nutrition, and growth. Here, we present a resource of organ-resolved, sex-specific, anatomically accurate models of newborn and infant metabolism, referred to as infant-whole-body models (infant-WBMs), spanning the first 180 days of life. These infant-WBMs were parameterised to represent the distinct metabolic characteristics of newborns and infants accurately. In particular, we adjusted the changes in organ weights, the energy requirements of brain development, heart function, and thermoregulation, as well as dietary requirements and energy requirements for physical activity. Subsequently, we validated the accuracy of the infant-WBMs by showing that the predicted neonatal and infant growth was consistent with the recommended growth by the World Health Organisation. We assessed the infant-WBMs’ reliability and capabilities for personalisation by simulating 10,000 newborn models, personalised with blood concentration measurements from newborn screening and birth weight. Moreover, we demonstrate that the models can accurately predict changes over time in known blood biomarkers in inherited metabolic diseases. By this, the infant-WBM resource can provide valuable insights into infant metabolism on an organ-resolved level and enable a holistic view of the metabolic processes occurring in infants, considering the unique energy and dietary requirements as well as growth patterns specific to this population. As such, the infant-WBM resource holds promise for personalised medicine, as the infant-WBMs could be a first step to digital metabolic twins for newborn and infant metabolism for personalised systematic simulations and treatment planning.

## Introduction

Infancy is a complex phase in human life, where numerous factors within an infant’s metabolism play together to enable rapid growth and healthy development of the body. Understanding the metabolism during infancy on an individual and population scale can be very beneficial, as the development in this early stage of life can have long-term consequences on human health.^1^

To comprehensively consider and understand the different factors influencing infant metabolism can pose challenges due to the complexity and scale of the system. Mathematical models provide a systematic approach to studying biological systems in detail, enabling hypothesis testing, variable adjustments, and efficient analysis of model outcomes. Various computational models have been developed to gain insights into infant development and disease processes. For instance, artificial neural network-based models have been developed to, e.g., predict neonatal metabolic bone disease,^2^ while mechanistic models have investigated the ratio of resting energy expenditure to body mass in childhood.^3^ These models can be adapted by integrating physiologic data and can be used to simulate complex metabolic processes, such as the dynamic coordination of macronutrient balance during infant growth,^4^ and to investigate infant skin permeability to topically applied substances.^5^ Additionally, physiology-based pharmacokinetic (PBPK) models have been developed to evaluate drug metabolism *in silico* for infants,^6^ which are aimed to complement the dose-extrapolation often performed for the determination of appropriate infant drug dosage.^7^

Given the complexity of whole-body metabolism in health and disease, genome-scale metabolic models (GEMs) also increasingly play a role in understanding human metabolism.^8–10^ GEMs are powerful tools to analyse complex metabolic networks as they enable the exploration of organism-level metabolic interactions.^11^ These models are assembled based on genomic, biochemical, and physiological data to generate a detailed description of molecular-level processes within a metabolic system.^12, 13^ They allow for an integration of various data sources, such as metabolomics,^14^ transcriptomic,^15^ proteomic,^16^ and metagenomic data.^17^ Importantly, using the constraint-based reconstruction and analysis (COBRA) approach, they can predict emergent metabolic properties. Numerous successive generic reconstructions of human metabolism have been published.^18–22^ These generic cell-level GEMs enable the investigation of human metabolism to further analyse metabolic functions and to elucidate the metabolic system and its relationship with diseases.^14, 23^ Recently, whole-body models (WBMs) of human metabolism have been introduced, which are sex-specific and organ-resolved.^8^ These WBMs have been developed starting from a generic, cell-level reconstruction,^21^ organ-specific information, and omics data to capture the metabolism of up to 26 organs and six blood cell types in an adult body.^8^ Each WBM contains over 80,000 biochemical reactions in an anatomically and physiologically consistent manner and can be parameterised with physiological, dietary, and metabolomic data. Importantly, these WBMs can also be personalised using clinical and omics data, e.g., metabolomic and metagenomic data,^8^ and thus represent a first step towards a virtual metabolic human, or digital metabolic twin. A first such digital metabolic twin has been recently generated and analysed for a Crohn’s disease patient over a time span of 16 months.^9^

The WBMs represent male and female adults at a resting stage, and therefore, are not suitable to investigate *in silico* infants, whose metabolism is optimised to support biomass accretion and growth. So far, infant metabolism has been modelled using generic cell-level metabolic models.^21, 24^ For instance, Nilsson et al^24^ developed the simulation toolbox for infant growth (STIG-met), which simulates the growth of an infant fed with breast milk and adjusted to the infant energy requirements based on body composition for maintenance and energy expenditure of major organs as well as muscular activity. This model has been used to predict the infant’s growth rate over the first six months of life in accordance with the recommendation by the World Health Organisation (WHO).^25^ However, this infant model does not allow for an organ-level analysis of the predicted fluxes, does not include energy demands due to, e.g., thermoregulation, cannot be personalised based on physiological and metabolomics data. Furthermore, the prediction of known biomarkers of inherited metabolic diseases (IMDs) with flux balance analysis (FBA)^26^ has been a recurring evaluation for cell-based models^20, 21, 27^ and organ-specific models^28^ as well as whole-body models^8^ based on the steady-state assumption but also in a time-dependent manner by applying dynamic parsimonious FBA.^29^

In this study, we present a resource of sex-specific, organ-resolved whole-body models of infant metabolism, infant-WBMs, spanning the first 180 days of life. We used detailed knowledge of infants’ physiology and metabolic processes as well as newborn screening data to develop personalised infant-WBMs. We accounted for organ-specific parameters and included in detail the energy demand for brain development, heart function, muscular activity, and thermoregulation over a period of six months. The resulting infant-WBMs could predict infant growth between zero and six months, based on their sex, body weight, measured metabolite concentrations, and further physiological parameters, and the predictions were in agreement with the growth recommendations by the WHO. Furthermore, the models could correctly predict changes in known blood biomarkers in IMDs on a day-to-day basis. As such, this resource represents an important step towards a better understanding of newborn and infant metabolism and will open novel avenues to investigate *in silico* infant metabolism in health and disease.

## Results

### Overview of the reconstruction process resulting in sex-specific, organ-resolved whole-body models of newborn and infant metabolism

This study introduces the sex-specific, organ-resolved whole-body models of newborn and infant metabolism (infant-WBMs), specifically designed for modelling newborns and infants up to the age of six months. The models were created using the adult sex-specific, organ-resolved WBMs^8^ as a starting point (Figure 1 (A)). Through an extensive literature review, we identified important metabolic, physiological, energetic, and nutritional differences between adult and infantile metabolism (Table S1, S2), and included them in the infant-WBMs as appropriate (Method section for details). For instance, an important difference is that infant organs grow, while most adult cell types, and thus organs, do not replicate. Hence, the infant-WBMs account for biomass growth reactions, summarising the required metabolites for cellular replication,^13, 30^ for all organs. Consequently, 259 and 236 reactions were added to the female and male infant-WBMs, respectively (Table S15). The difference in reaction numbers was due to the difference in sex-specific organs. Additionally, the infant-WBMs present organ weights and blood flow supply rates corresponding to the different ages of the infants based on *in vivo* data.^6^ Other infant-specific physiological parameters include the heart rate, stroke volume, cardiac output, hematocrit value, creatinine concentration in urine, and glomerular filtration rate (Table S2). Moreover, the excretion of water and other metabolites were adapted such that the urine flow rate and perspiration (skin) as well as excretion of water through the air, faeces, and urine resembled infant metabolism (Method section 1.5). Organ-specific energy requirements are quite different between infants and adults^31^ and had to be adapted to represent the requirements for healthy infant development. For instance, for the proper development of the brain, we implemented these energy requirements by adjusting the lower bound on the brain ATP reaction (VMH ID: Brain DM atp c), which monotonously increases in accordance with the age of the infant-WBM (Method section 1.7). Similarly, the heart energy requirement is infant-specific, depending on the infant’s heart weight (Method section 1.9). With increasing age, activity-based energy requirements also increase, which we accounted for by adding constraints on the muscle ATP demand reaction (VMH ID: Muscle DM atp c) in accordance with a published activity model^24^ (Method section 1.10). Another important infant-specific energy demand is thermoregulation, which maintains the infant’s body temperature, as infants cannot shiver before the age of six months.^32, 33^ We implemented the energy requirement for the nonshivering thermogenesis process in the adipose tissue (Method section 1.8). We also applied infant-specific nutritional constraints to represent breastfed infants with a daily increasing diet corresponding to the infant’s growth.^24^ Note that we assumed the infants to be exclusively breastfed until the age of six months, as recommended by the WHO.^34^ The diet composition was based on the human milk decomposition at the Virtual Metabolic Human database^35^ (VMH, https://www.vmh.life/) and adapted where necessary (Method section 1.6). In general, we assumed that the composition of breast milk remains constant during the initial six months. However, breast milk composition is very dynamic and varies over time according to the needs of a growing child,^36^ which might be the reason that we had to adjust some metabolic components within the diet throughout the six-month evaluation (see Method section). Finally, we used measured blood metabolite concentrations from newborn screening of 10,000 newborns to adjust the metabolite concentration ranges for 17 amino acids and 12 acylcarnitines (Table S3) (Method section 2).

**Figure 1:**
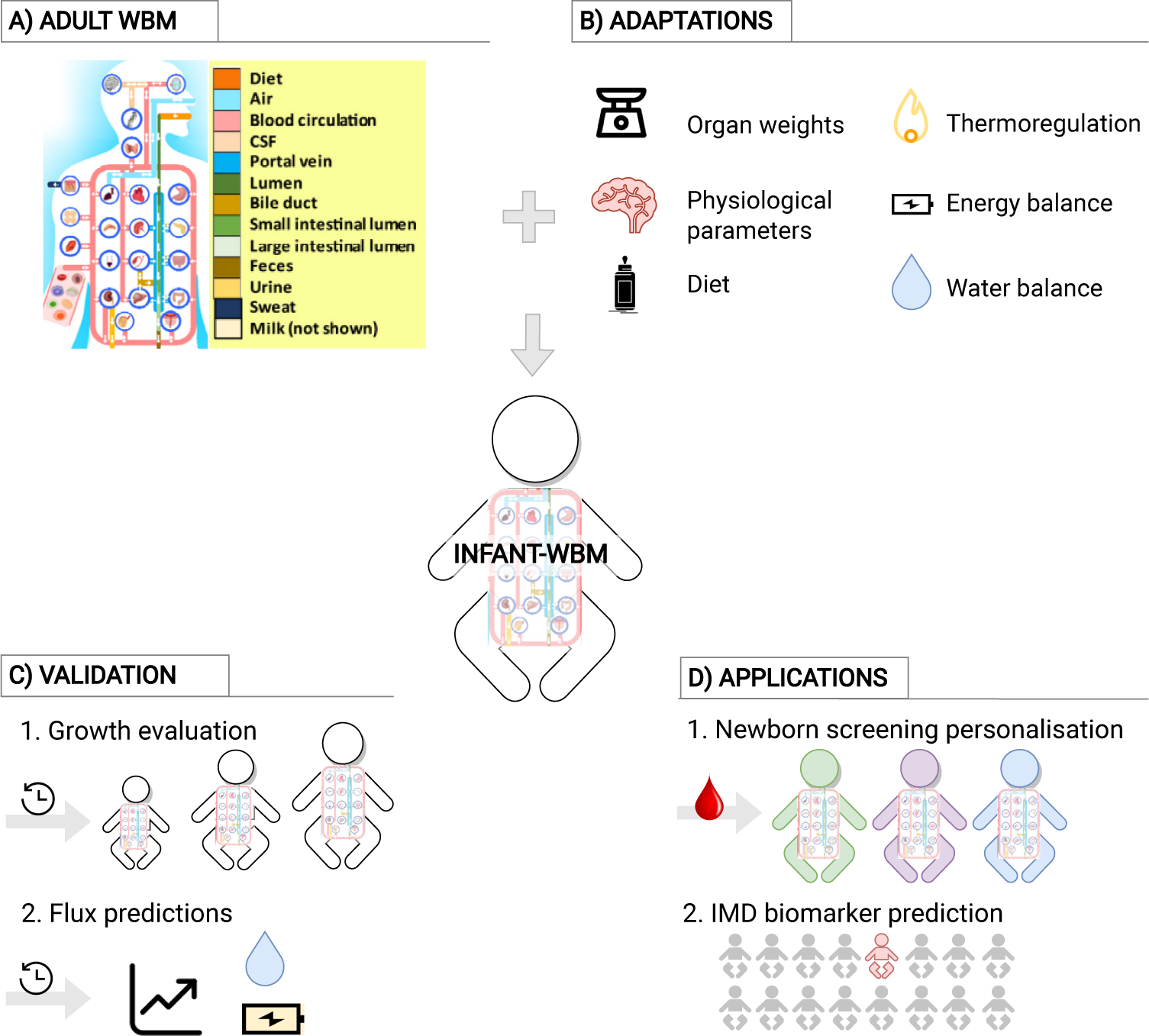
Overview of the reconstruction process and sample applications of the infant-WBMs. (A) The infant-WBMs were derived from the adult WBMs, figure taken from Thiele et al.^8^ (B) Main adaptions made to the adult to generate the infant WBMs, including the organ weights, physiological parameters, diet intake, thermoregulation, energy balance, and water balance. (C) Validation of the infant-WBMs with growth evaluation and flux predictions. (D) Application of the infant-WBMs for newborn screening personalisation and biomarker prediction of inherited metabolic diseases (IMDs).

Overall, the female infant-WBM accounts for 1,724 unique genes (2,071 transcripts), 85,662 reactions, and 60,436 metabolites, whereas the male infant-WBM accounts for 1,724 unique genes (2,071 transcripts), 83,149 reactions, and 57,980 metabolites. Taken together, the sex-specific infant-WBMs represent comprehensively metabolic, physiological, energetic, and nutritional features of infants aged between zero and six months.

### Validation - Prediction of growth rate over time

A critical step in modelling biological systems is the validation process and comparison with the actual organism, in our case, the human infant. An important difference between the metabolic models of adults and infants lies in the capacity of infants to undergo growth, leading to increases in total body weight and individual organ weights. Hence, to validate the infant-WBMs, we computed the growth curves for each sex and compared them with the recommendations of the WHO^25^ (Figure 2 (A), (B), Methods). Therefore, we used FBA^26^ to maximise the flux through the whole-body biomass reaction, which accounts for the contribution of each organ’s biomass growth reaction weighted based on the relative organ weight. On each successive day, the body weight was adjusted based on the previous day’s growth rate, followed by updates to the organ weights according to the adjusted body weight. The initial newborn weights (day 0) were set to be 3,300 g and 3,200 g for the male and female infant-WBMs, respectively, and reaching a weight of 7,900 g and 6,090 g at day 180 (Figure 2 (A), (B), Table S4). This weight gain corresponded to a daily average of 0.49% and 0.43% of gain in body weight for male and female infant-WBMs. Overall, the calculated infant weights for each day of life during the first 180 days were in very good agreement with the WHO recommendations, as they were within the WHO quartiles (Figure 2 (A), (B), Table S4), and they were also consistent with previous growth predictions performed with the generic genome-scale models of human metabolism.^21, 24^ We conclude that the iterative adaptation of body weight determined by the growth rate and the dynamical adjustment of the ATP demands for the brain, heart, thermoregulation, muscular activity, and milk intake according to the age of the infant resulted in an accurate growth trajectory. In the following sections, we demonstrate two further validation steps and two applications of the infant-WBMs.

**Figure 2:**
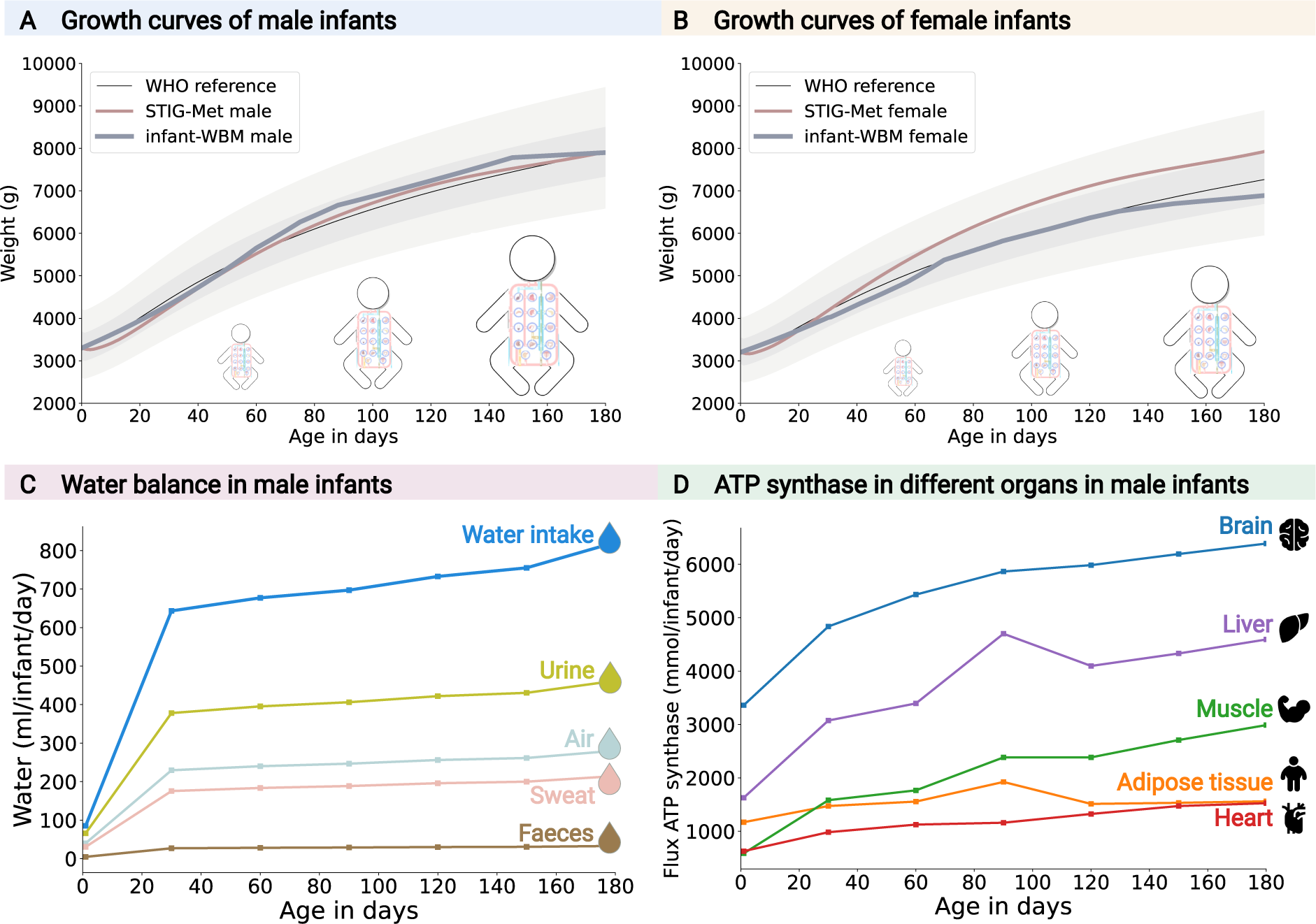
Validation of infant-WBMs over the first six months of *in silico* life. (A) Male and (B) female infant-WBM growth predictions in comparison with WHO quartiles^25^ and predictions by the STIG-Met model.^24^ A flux value of 1.01 through the whole-body biomass reaction corresponds to a 1% gain in biomass, i.e., an increase in the body weight of the infant by 1%. For the validation, the male infant models start on day zero with a body weight of 3, 300 g (A) and the female infant-WBMs with a body weight of 3, 200 g (B). (C) Water intake and excretion fluxes in male infants computed over the first six months. (D) Flux through the ATP synthase reactions in the brain, muscle, adipose tissue, liver, and heart in male infants over the first six months.

### Validation - Prediction of metabolic fluxes within the infant body

The infant-WBMs enable a comprehensive assessment of metabolic fluxes at the organ, sub-system, and reaction level in infants. Here, we discuss the validation results of the male infant-WBM. The data and results for the female infant-WBM can be found in the supplementary material (Table S7 (A)-(B), Supplementary Figure S1).

#### Water balance flux predictions

The water balance describes the equilibrium between water intake and water loss within the human body. For both adults and infants, it is crucial to maintain proper hydration and to ensure the normal functioning of various physiological processes.^37^ Correspondingly, the dietary intake of water determines most of the excretions through urine, faeces, skin, and respiration. In addition, metabolic water is produced inside a living organism as an end product of the oxidation of energy-containing substances in their food.^38^ Here, we determined whether the infant-WBMs could correctly predict water loss. Therefore, we computed the flux distribution through each infant-WBM (Method section). As expected, and consistent with the applied constraints (Table S12 (B)), most of the water was excreted via the urine (Figure 2 (C)). We predicted urine excretion to be between 66 ml per day on day 1 and increasing to 462 ml per day on day 180 (Table S6(A)). These predicted values were, in general, comparable with measurements in infants, where urine excretion of infants up to one year has been approximated as 2· weight(kg) ·24 (ml/day),^39^ thus ranging from 158 - 379 ml per day between ages 0 – 6 months, assuming a body weight of 3.3 to 7.9 kg. Water loss through air and sweat represented the next highest values, and the predictions of 71 - 495 ml per day (Figure 2 (C), Table S6 (A)) were consistent with estimations on water evaporation through skin and air being 148.5 - 355.5 ml per day for 0 - 6 months.^40^ The predicted faecal water loss ranged between 4.8 ml/day and 33 ml/day, which is comparable to the reported 5 ml/kg per day (i.e., 16.5 ml/day and 39.5 ml/day for 0 - 6 months) as water loss via defaecation for newborns.^40^ Overall, the predicted water excretion ranged from 141 ml per day on day 1 to 990 ml/day on day 180, with an average of 783 ml per day across all predicted time points (Table S6 (A)). This predicted average value compared well with the reported mean water excretion of 900 ml/day from 78 male babies aged between 8 and 180 days (mean=36 days).^38^ Notably, the predicted water excretion on day 30 was 810 ml/day. It is important to note that these predicted output values also accounted for the metabolic production of water, which was between 56 ml/day and 169 ml/day (Table S6 (A)), highlighting that overall the infant-WBMs realistically captured water production and consumption. While our predictions did not perfectly match the reported values, and we had applied constraints on the excretion reactions, this example demonstrates that the infants-WBMs can capture the known water balance. This capability is an emergent feature of the infant-WBMs as they have not been trained to recapitulate the water balance.

#### Prediction of ATP synthase in different organs

The energy balance of the infant-WBM describes the intake and use of energy within the infant’s body. Unlike generic genome-scale models of human metabolism, such as Recon3D^21^ and Human1,^22^ the infant-WBM allows the allocation of distinct energy demands to specific organs for their respective functions. Consequently, we can predict different aspects of an infant’s energy balance, taking into consideration the energy intake from the breast milk diet and the demands of different organs required for brain development, thermoregulation, heart function, and physical activity. To represent the organ-specific energetic requirements, the infant-WBMs contain ATP demand reactions, which represent the ATP hydrolysis for non-growth-associated metabolic processes (e.g., muscle or brain activity). As the applied constraints through these reactions increase, the flux through the ATP synthase reaction, which is part of the oxidative phosphorylation in these organs, was expected to increase accordingly. Consistently, the brain ATP synthase had the highest predicted flux value being 3,360 mmol/day/infant on day 1 and 6,388 mmol/day/infant on day 180, followed by the liver ATP synthase (Figure 2 (D), Table S6 (B)).

Next, we examined the temporal changes in ATP synthase flux values in the brain, muscle, liver, adipose tissue, and heart, to shed light on the dynamic energy metabolism during early infancy (Figure 2 (D)). The highest increase in ATP synthase flux was predicted for the muscle, which was five times higher on day 180 compared to day 1. This increase was a direct reflection of the increase in the lower bound on the muscle ATP demand (Figure 5), which represents the increase in physical activity of the growing infant. In contrast, the adipocyte ATP synthase flux increased only by 33.5%. Interestingly, the predicted liver ATP synthase flux increased from 1,629 mmol/day/infant on day 1 to 4,591 mmol/day/infant on day 180, representing an increase of 2.8 times. No constraints were placed on the liver ATP demand reaction and thus, this predicted increase was an emergent feature of the infant-WBMs representing the increase in overall metabolic activity during infant growth.

When correcting for the infant’s weight, the ATP synthase flux from the brain and adipose tissue decreased from day 1 to day 180 (from 1,018 mmol/kg/day to 809 mmol/kg/day, and from 355 mmol/kg/day to 198 mmol/kg/day, respectively). The heart ATP synthase flux was predicted to be nearly unchanged during the infant’s growth (190 mmol/kg/day on day 1 vs 193 mmol/kg/day on day 180). In contrast, liver ATP synthase flux increased by 18% (494 mmol/kg/day to 581 mmol/kg/day) and the muscle ATP synthase flux doubled (178 mmol/kg/day to 378 mmol/kg/day). For the muscle, we also investigated the ATP produced from glycolysis, which remained nearly unchanged, when corrected for weight, from day 1 to day 180 (149 mmol/kg/day vs 150 mmol/kg/day). This result shows that, *in silico*, the additional energy requirement for physical activity was covered by an increase in oxidative phosphorylation. Consistently, the Cori cycle was active in the infant-WBMs with the highest gluconeogenesis flux, determined as the flux through the liver glucose-6-phosphate phosphatase (VMH ID: Liver G6PPer), being highest on day 60 (i.e., 105 mmol/kg/day) when corrected for weight (Table S6). This result was consistent with reports that in rats, the highest glucose-6-phosphate phosphatase was found at age 7 days^41^ corresponding to 60.2 human days, assuming that during the weaning phase, a rat day corresponds to 8.6 human days.^42^ Taken together, this example demonstrates that the infant-WBMs recapitulate known ATP requirements and metabolism.

### Application - infant-WBMs enable integrative analysis of metabolome data from 10,000 healthy newborns

Newborn screening programs worldwide aim at early, ideally, presymptomatic identification of treatable rare diseases, such as IMDs, to reduce morbidity and mortality.^43^ For this aim, metabolic, enzymatic, and genetic parameters are analysed with dried blood samples taken in the first days of life (i.e., in Germany at 36-72 h of life). Most metabolic parameters are analysed by tandem mass spectrometry to quantify their concentration. This process generates extensive individualised metabolomic data. However, the specific conditions examined and the number of metabolites measured in each newborn screening program vary between countries.^43, 44^ Here, we generated personalised infant-WBMs using anonymised data obtained from the newborn screening laboratory at Heidelberg University Hospital (UKHD), Germany. The full newborn screening data set comprised information from over 2 million individuals and encompassed 29 amino acids and acylcarnitines, which could be mapped to the metabolites present in the blood compartment of the infant-WBMs (Table S3). The metabolomic data obtained from newborns provides a snapshot of metabolite concentrations in the blood at a specific time point. These metabolite concentrations can be incorporated into the infant-WBMs by updating the bounds on the uptake fluxes from the blood compartment into individual organs (Method section 2). Additionally, the birth weight and sex of the newborns were used to adjust all weight and sex-specific parameters within the infant-WBMs. This adaptation ensured that the personalised infant-WBMs accurately reflected the individual characteristics of each newborn. From the full newborn screening data set, we evaluated a uniformly random sampled subset of 10, 000 male newborn models with gestational age greater or equal to 38 weeks (further details in Method section 2). The mean measured birth weight in the subset was 3, 508 g, ranging from 1, 770 g to 5, 590 g (Figure 3 (A)). We then computed the *in silico* growth rates using FBA. From all 10,000 personalised infant-WBMs, 8,736 (87.36%) models had a growth rate between 1.0089 - 1.0092 (mean = 1.0091 ± 0.00008), corresponding to a weight gain of 0.89 - 0.92% per day (Figure 3 (B), Table S8). Interestingly, from the remaining models 1,108 models (11.08%) had a daily growth rate between 0.22 and 0.996 (mean = 0.74 ± 0.23) implying that the infants would lose weight (Figure 3 (B), Table S8). Loss of birth weight up to 10% in the first few days of life is normal,^45^ and presumingly, due to fluid loss through urination.^46^ It is to be noted that we did not change any water-related constraints in the personalised infant-WBMs. Unfortunately, no further information on the infants’ follow-up weight (e.g., on day 3), general health status (besides the tested IMDs), potential feeding problems, delivery mode, or the age of the mother where available, to validate the predicted loss in weight. However, all these factors have been associated with excessive weight loss in newborns.^47^

**Figure 3:**
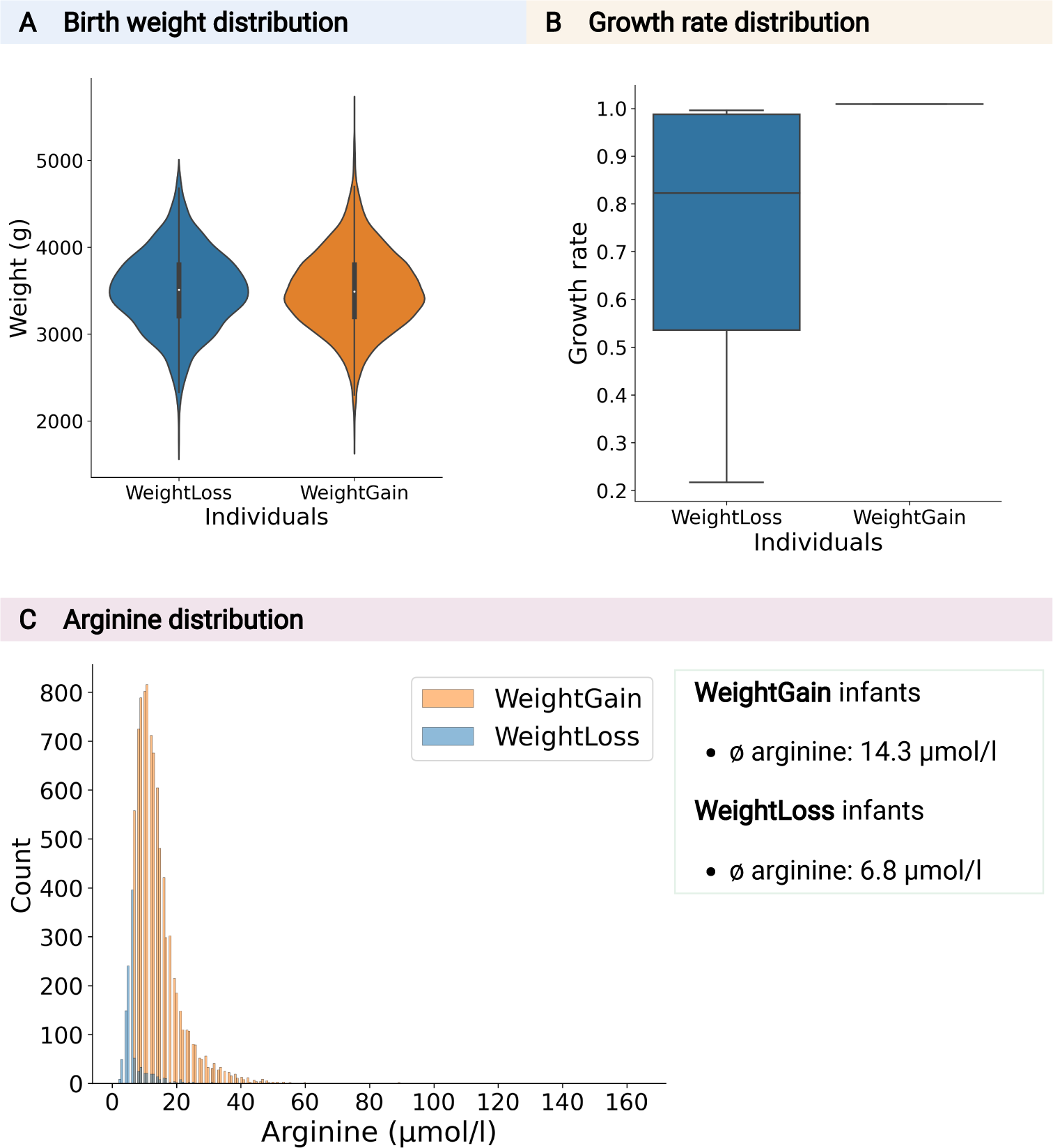
Evaluation results of 10,000 male infant-WBMs. (A) Birth weight and (B) growth rate distribution of infants with predicted weight loss (WeightLoss) and infants with predicted weight gain (WeightGain). (C) Comparison of measured arginine concentration in infants.

To further shed light on the predicted weight loss, we investigated potential metabolic reasons. We did not observe any correlation between the birth weight and the predicted growth rate, as the birth weight was similarly distributed between the weight gainers and weight losers (Figure 3 (A)). We also could not find any correlation with gestational age (Supplementary Figure S2). When we compared the distributions of all the personalisation variables, we found that the measured blood concentration of arginine was very low in weight losers (mean = 6.8 ± 4.1µmol/l) compared to the weight gainers (mean = 14.3 ± 7.4µmol/l; Figure 3 (C)). The distribution of the arginine concentrations of the weight gainer and weight loser infants was significantly different (Wilcoxon rank sum test, *p* = 1.8*e^−^*^227^). Increasing the blood arginine concentration *in silico* to 14 µmol/l enabled 762 (69%) of the weight loser infant-WBMs to grow at rates of at least 1.005, confirming blood arginine concentrations as one of the growth-limiting variables in these infant-WBMs. For the remaining 346 (31%) of the weight loser infant-WBMs, we could not find any single measured blood metabolite that could explain the predicted weight loss. We then tested whether dietary supplementation could also achieve higher *in silico* growth rates in the weight losers using a dedicated algorithm, the nutrition algorithm^48^ (Figure 3), but we could not find any alternative. Note that in the way the diet algorithm is designed, it does not change the blood concentrations in the infant-WBMs, but rather asks whether there is any metabolite that could substitute for the arginine limitation or any other limitation. Arginine is known to be an essential amino acid in newborns and infants,^49^ while not being an essential dietary amino acid in healthy adults,^50^ and our *in silico* modelling confirmed this. Decreased plasma l-arginine levels in organic acidurias have been suggested as a potential cause of growth retardation in children and adolescent patients with methylmalonic acidemia and propionic acidemia.^51^ Moreover, a study of Danish schoolchildren reported an association between dietary arginine intake and growth rate.^52^ For the reason of low blood arginine levels in our infants, we can also only speculate. Diet is a potential source of variation. However, breast milk has been reported to be a poor source of arginine, while its metabolic precursors are highly abundant.^53^ Mutations in the genes associated with arginine synthesis could be another source, thereby limiting the production of arginine from metabolic precursors in the small intestine.^49^ Finally, differences in gut microbial composition may also contribute to variations in the blood concentration of arginine and its precursors. Accordingly, we found that gut microbiomes from healthy infants were enriched in reactions associated with “Arginine and proline metabolism” compared with healthy adults (average 13.5% ± 2.9% vs 10.7% ± 1.2%)^54^ (Table S13). Taken together, this application demonstrates how the infant-WBMs can be personalised based on newborn screening metabolomic data to investigate emerging metabolic properties.

### Application - infant-WBMs accurately predict known biomarkers of inherited metabolic diseases

Single gene defects in biochemical pathways can result in IMDs.^56^ While IMDs are individually rare (from 1:1,000,000 to 1:10,000 newborns), the cumulative incidence of the over 1,880 recorded IMD,^56^ associated with over 1,367 unique genes,^55, 57^ is high (ranging from 1:2,500 to 1:800 newborns).^58^ The infant-WBMs cover 595 of the 1,364 IMD genes (43%) and are associated with 636 IMDs, providing ample opportunity to investigate known and novel biomarkers (Table S16). These 595 IMD-associated genes are distributed over all 23 disorder classes of IMDs as defined by Ferreira et al^55^ (Table 1).

**Table 1:** IMD groups/classes with their associated number of diseases and coverage in the infant-WBM model. *The classification was based on.^55^

| IMD groups/classes* | Diseases<br>per<br>class | Infant-WBM<br>associated<br>diseases per<br>class | Fraction |
| --- | --- | --- | --- |
| 1 - Disorders of amino acid metabolism | 113 | 91 | 0.805 |
| 2 - Disorders of peptide and amine metabolism | 20 | 14 | 0.700 |
| 3 - Disorders of carbohydrate metabolism | 67 | 51 | 0.761 |
| 4 - Disorders of fatty acid and ketone body metabolism | 28 | 26 | 0.929 |
| 5 - Disorders of energy substrate metabolism | 28 | 26 | 0.929 |
| 6 - mtDNA-related disorders | 40 | 11 | 0.275 |
| 7 - Nuclear-encoded disorders of oxidative phosphorylation | 88 | 47 | 0.534 |
| 8 - Disorders of mitochondrial cofactor biosynthesis | 28 | 3 | 0.107 |
| 9 - Disorders of mitochondrial DNA maintenance and replication | 17 | 5 | 0.294 |
| 10 - Disorders of mitochondrial gene expression | 65 | 1 | 0.015 |
| 11 - Other disorders of mitochondrial function | 53 | 14 | 0.264 |
| 12 - Other disorders of intermediary metabolism | 9 | 6 | 0.667 |
| 13 - Disorders of lipid metabolism | 145 | 94 | 0.648 |
| 14 - Disorders of lipoprotein metabolism | 30 | 7 | 0.233 |
| 15 - Disorders of nucleobase, nucleotide and nucleic acid metabolism | 151 | 38 | 0.252 |
| 16 - Disorders of tetrapyrrole metabolism | 19 | 17 | 0.895 |
| 17 - Congenital disorders of glycosylation | 137 | 61 | 0.445 |
| 18 - Disorders of organelle biogenesis, dynamics and interactions | 111 | 3 | 0.027 |
| 19 - Disorders of complex molecule degradation | 73 | 36 | 0.493 |
| 20 - Disorders of vitamin and cofactor metabolism | 71 | 40 | 0.563 |
| 21 - Disorders of trace elements and metals | 34 | 8 | 0.235 |
| 22 - Neurotransmitter disorders | 67 | 13 | 0.209 |
| 23 - Endocrine metabolic disorders | 50 | 23 | 0.460 |
| Total | 1,444 | 636 | 0.44 |

The highest coverage of disorder classes was for “Disorders of fatty acid and ketone body metabolism” and “Disorders of energy substrate metabolism”, where the infant-WBMs cover 26/28 IMDs in the respective classes. Also, a high number of IMDs of “Disorders of lipid metabolism” (94/145 IMDs) and “Disorders of amino acid metabolism” (91/113 IMDs) are accounted for. The lowest coverage was achieved for “Disorders of mitochondrial gene expression” (1/65 IMDs) and “Disorders of organelle biogenesis, dynamics and interactions” (3/111). This low coverage was not surprising as the associated pathways are traditionally not covered in metabolic reconstructions. Furthermore, the 596 genes correspond to 14,187/85,662 reactions (16.6%) in the female infant-WBMs and 13,633/83.149 reactions (16.4%) in the male infant-WBMs. This mapping provides a prime opportunity to expand the infant-WBMs in future efforts and a starting point for investigating IMDs and emergent phenotypes *in silico*.

Since the publication of the first human metabolic reconstruction,^18^ the prediction of known biomarkers of IMDs^27^ has been a recurring evaluation for cell-based^20, 21^ and whole-body models^8^ as well as models of organ-specific metabolism.^28, 29^ Hence, to further demonstrate the predictive capacity of the infant-WBM models, we first analysed the biomarker prediction for phenylketonuria (PKU) (OMIM: #261600), which is an inborn error of phenylalanine metabolism (synthesis of tyrosine from phenylalanine by the phenylalaninhydroxylase) and is, if untreated, associated with global developmental delay and severe intellectual impairment of patients.^59^ Using an established method to predict biomarkers for IMDs *in silico*^8^ (Method section 3.5), we predicted for the male and female infant-WBMs the known PKU biomarkers as well as 27 further metabolites routinely measured in the dried blood spots for newborn screening at the UKHD (Figure 4, Table S3). We compared the prediction of the wild-type (healthy) model with the knock-out (PKU) model after deleting the corresponding reactions for tetrahydrobiopterin: oxygen oxidoreductase (VMHID: r0399, PHETHPTOX2) in all organs having the known defective gene phenylalanine hydroxylase (VMHID: 5053.1) and calculating the relative flux change *f* (section 3.5). In both female and male infant-WBMs, the phenylalanine flux was predicted to increase more than 300% and the tyrosine flux was predicted to decrease by 91% in female and 97% in male models. Thus, the known biomarkers phenylalanine and tyrosine showed by far the highest relative change in flux prediction, whereas the flux through all other newborn screening metabolite reactions showed no or small relative flux changes below 3% (Figure 4). Interestingly, 12 of the 29 metabolite fluxes (alanine, arginine, argininosuccinate, aspartate, citrulline, glutamic acid, glycine, homocitrulline, methionine, phenylalanine, proline, tyrosine) were predicted to change in both female and male models as well as two metabolite fluxes (histidine, isovalerylcarnitine) that were only predicted to change in the female model and one metabolite flux (glutarylcarnitine) that was only predicted to change in the male model (Figure 4).

**Figure 4:**
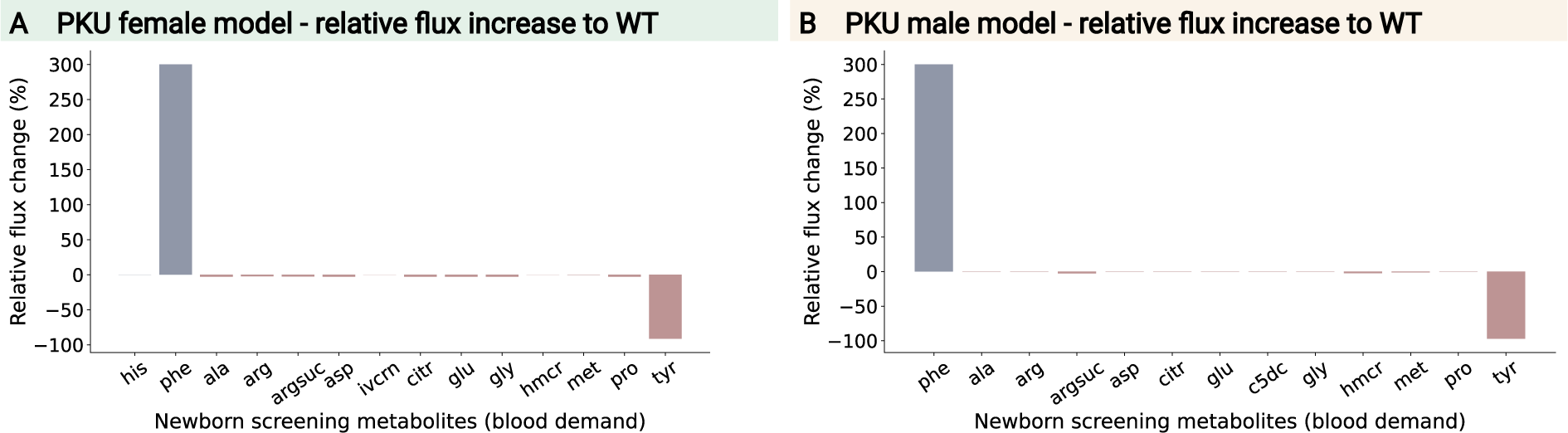
Relative change of flux through blood demand reactions of metabolites measured in newborn screening for a female (A) and male (B) infant-WBM comparing the wild type with a PKU knock-out model. Abbreviations in VMH IDs: ala - alanine, arg - arginine, argsuc - argininosuccinate, asp - aspartate, c5dc - glutarylcarnitine, citr - citrulline, glu - glutamic acid, gly - glycine, his - histidine, hmcr - homocitrulline, ivcrn - isovalerylcarnitine, met - methionine, phe - phenylalanine, pro - proline, tyr - tyrosine.

To further demonstrate the predictive capacity of the infant-WBM models, we chose three IMDs, which are part of the German national newborn screening, including PKU, isovaleric aciduria (IVA) (OMIM: #243500), and glutaric aciduria (GA1) (OMIM: #230800),^60^ since in newborn screening dried blood samples are analysed regarding significant increases or decreases of concentrations of the IMD specific biomarkers. Additionally, we also investigated Gaucher disease (GD) (OMIM: #230800) and kynureninase deficiency (KD) (OMIM: #236800) (Table 2, Table S11 (A)). For each IMD, we predicted and evaluated the flux through their known biomarkers over the time course of the first six months of a female infant’s life (Table S11 (B)). The data and results for the biomarker prediction of the male infant-WBM can be found in the supplementary material (Figure S2, Table S11 (C)). We evaluated the predicted fluxes qualitatively by comparing the flux through the known biomarker reactions in the wild type (healthy) and IMD (disease) model and noted whether an elevation or reduction of the flux was predicted (Table 2). For all IEMs and all the biomarker, the predictions were consistent over all three time points and consistent with the change in biomarkers reported in IEMBase.^56^

**Table 2:** Qualitative flux change through biomarker reactions in blood on day 1, 90, and 180 for female infant-WBM and for female adult WBM,^8^ when maximising the respective biomarker reactions in the knockout (disease) and the wild type (healthy) model. A predicted flux increase is highlighted with an up-arrow ⇑ and a decrease with a down-arrow ⇓. PKU - phenylketonuria (OMIM: #261600), IVA - isovaleric aciduria (OMIM: #243500), GA1 - glutaric aciduria (OMIM: #230800), GD: Gaucher disease (OMIM: #230800), KD - Kynureninase deficiency (OMIM: #236800). Phe - phenylalanine, tyr - tyrosine, 4hphac - p- hydroxyphenylacetic acid, phpyr - phenylpyruvic acid, ivcrn - isovaleryl carnitine, c5dc - glu- taryl carnitine, gluside - D-glucosyl-N-acylsphingosine, C02470 - xanthurenic acid, kynate - kynurenic acid. All metabolite abbreviations are given as VMH IDs.

| Diseases | Metabolites | Day 1 | Day 90 | Day 180 | Adult |
| --- | --- | --- | --- | --- | --- |
| PKU | phe | ↑ | ↑ | ↑ | ↑ |
| | tyr | ↓ | ↓ | ↓ | ↓ |
| | 4hphac | ↓ | ↓ | ↓ | ↑ |
| | phpyr | ↑ | ↑ | ↑ | ↑ |
| IVA | ivcrn | ↑ | ↑ | ↑ | ↑ |
| GA1 | c5dc | ↑ | ↑ | ↑ | ↑ |
| GD | gluside | ↑ | ↑ | ↑ | ↑ |
| KD | C02470 | ↑ | ↑ | ↑ | ↑ |
| | kynate | ↑ | ↑ | ↑ | ↑ |

Overall, this analysis demonstrates that the infant-WBMs have the ability to predict known biomarkers related to a specific IMD over a time frame of six months evaluated on three time points, day 1, day 90, and day 180. Moreover, all biomarker changes were also consistent with changes predicted by the adult WBMs, except for the prediction of p-hydroxyphenylacetic acid for PKU (Table 2). For the adult WBMs, it has been already shown that they have a good predictive capacity for IMDs,^8^ where in a simulation of predicting biomarkers in blood, urine, or cerebrospinal fluid (CSF), 85.3% of biofluids have been qualitatively predicted correctly by the female adult WBM and 84.9% by the male adult WBM.^8^ Taken together, we showed that the infant-WBMs could correctly predict biomarkers for a range of metabolites and IMDs over the first six months of life, which may be of value for newborn screening but also for biochemical tests performed on patients with suspected IMDs.

## Discussion

Here, we presented a resource of infant-WBMs, i.e., genome-scale metabolic sex-specific whole-body models for newborns and infants, which incorporate detailed knowledge of infants’ physiology and metabolic processes as well as newborn screening data for personalisation over the first six months of life. By including organ-specific parameters and detailed information on the energy demand for brain development, heart function, muscular activity, and thermoregulation, we could successfully model the newborn and infant metabolism over a period of six months. We showed that when maximising the whole-body biomass reaction, we could accurately predict the infant’s growth during this time span, in accordance with growth recommendation from the WHO,^25^ for both male and female infants (Figure 2 (A), (B)). Moreover, we validated the model by analysing the flux prediction of whole-body metabolism. For instance, we confirmed that the amount of daily water excretion and interorgan energy metabolism within the infant-WBMs (Figure 2 (C), (D)) agreed with *in vivo* measurements of infants within a reasonable range throughout the six-month time frame. Furthermore, we used the models in two applications related to newborn screening. First, we evaluated the robustness of the models by creating 10,000 personalised infant-WBMs utilising the sex, birth weight, and 29 metabolite concentrations measured during newborn screening (Figure 3). Second, we showed that the infant-WBMs are able to correctly predict known metabolic biomarkers for five IMDs at three different time points (Table 2). This analysis could be a new way of investigating IMDs as the harmful accumulation of metabolites, such as the mitochondrial accumulation of isovaleryl-CoA in IVA patients, is a key problem.^61^ For many diseases, such as Gaucher disease, investigations of metabolic flux are important as the alteration of flux is central to the management of the disease.^62^ In particular, in newborn screening, the development of the newborn’s metabolism is essential, as it allows early, ideally, presymptomatic identification of treatable IMDs, enabling beneficial treatment of the affected newborns.

### Limitations of the infant-WBMs

A mathematical model of a complex biological system, such as infant metabolism, will never perfectly resemble the underlying biological system, which is due to necessary assumptions and simplifications. Hence, the infant-WBMs also have limitations based on decisions taken during the development process. The infant-WBMs were derived from the female and male adult WBMs, and the contained reactions were only extended for the biomass growth reactions in every organ, which is an important difference between adult and infant metabolism. Additionally, infant-specific constraints were placed on the organ’s energy demands, diet, and physiological parameters. However, no further changes were made to the reaction content or applied constraints, although the metabolic profile varies at different stages of human life.^63^ For instance, the analysis of urine samples showed that the activity of the pathways associated with amino acid metabolism is significantly different between infants aged six months and one year.^63^ Hence, adding further constraints to the involved reactions depending on age could further improve the accuracy of the models’ predictive capacity of infant metabolism.

Furthermore, the infant diet is a very important and sensitive part of the model. Here, we assumed that the infants are exclusively breastfed, which is recommended by the WHO.^34^ However, a study on breastfeeding among U.S. children showed that, in 2019, 35.3% of all children were supplemented with infant formula before six months of age.^64^ Hence, in a future study, different diets for infants, e.g., including formula, could be considered. By this, also the dietary effects on the infant metabolism could be analysed and using the nutrition algorithm^48^ missing diet components could be predicted in a personalised manner. We further assumed that, except for some metabolites, the milk composition over the first six months does not change, although the breast milk composition can be very dynamic and varies over time according to the needs of a growing child.^36^ However, to integrate these dynamic changes within the model, detailed measurements of the diet composition at several time points per day would be necessary over the six-month time frame. Still, this would only represent a personalised diet for a specific infant as the diet composition can vary depending on environmental pollution,^65^ exogenous chemicals, such as drugs and synthetic compounds,^66^ and the mother.^67^ The mother influences the milk composition by her nutrition, which has been reported to impact concentrations of fatty acids and fat- and water-soluble vitamins in the milk^68^ as well as by her milk production quantities, which impacts the concentration of fat, protein, and lactose in the milk.^67^ The assumed constant milk composition over time may be a reason for the adjustments that had to be performed to some metabolic components within the diet throughout the six-month evaluation to ensure the model’s feasibility. In particular, the dietary intake of L-lysine had to be decreased and L-cysteine had to be increased for the model to be able to predict a reasonable growth rate. Overall, we found some of the predictions sensitive to the diet input, thus highlighting the need for proper constraints. It should also be noted that we did not include the gut microbiome, which is known to provide essential macro- and micronutrients to the host metabolism,^69^ and thus can complement the dietary inputs. Moreover, the infant-WBMs do not account for protein turnover, which is age-dependent and important for growth. However, in order to represent the protein turnover correctly, we would have to include the synthesis and degradation of the major proteins, which has been done for microbes (e.g.,^70, 71^), but was out of scope for this study.

### Future and implications

The integration of microbiome data into infant-WBMs represents a natural progression, as it has already been part of the adult WBMs.^8^ This integration would also enable the consideration of the mode of birth, as studies have shown that the gut microbiota composition of C-section delivery newborns shows a microbiome that closely resembles that found in the environment and the mother’s skin, whereas vaginally delivered newborns have a microbiome more similar to the vaginal microbiome.^72^

Furthermore, by incorporating different diets for infants, the modelled age range of the children could be expanded. This extension would also require additional changes in physiological information within the models, such as changes in the organ weights at different stages. Ultimately, it would be possible to develop WBMs that encompass all early stages of human life, starting from newborns and progressing to the existing adult models. This comprehensive coverage of different life stages would provide valuable insights into the dynamic nature of human metabolism throughout the lifespan.

The personalisation of the infant-WBMs, which we showed on a subset of 10,000 newborns, allowed for incorporating personalised parameters and contextualising the models within the infant population. This approach provides valuable insights into the individualised metabolic characteristics of infants, paving the way for personalised interventions and an improved understanding of early-life metabolism. The infant-WBMs enable newborn screening centres world-wide to use their own measured metabolite concentrations to create personalised metabolic models of their patients. These personalised models could be eventually used to test therapies and treatments *in silico*, as, due to the extreme variability of IMDs, their management and therapy have to be personalised for each patient, based on the patient’s diagnosis and phenotype.^62^ In the era of precision medicine, the ability to accurately predict an infant’s metabolic response to various dietary interventions holds immense potential for personalised nutritional strategies, clinical decision-making, and improving the management of IMDs in infants by analysing how a phenotype emerges given a primary mutation on the background of their whole personal genome.^73^

Personalised WBMs could also be used to analyse the metabolic impact of exogenous chemicals, such as drugs and synthetic compounds, that may be transferred from human milk to infants.^66^ For instance, the personalised infant-WBMs could support newborn screening for IVA, which is suffering from an increasing number of false-positive screening results, due to antibiotics given to pregnant women to treat urinary tract infections^74^ and pivalate-containing creams.^75^ Moreover, drug dosage determination for children is very important as their immature drug metabolism is often associated with drug toxicity^76^ and the pharmacokinetics and pharmacodynamics of drugs are often different in children and adults.^77, 78^ However, almost 50% of prescription drugs lack age-appropriate dosing guidelines^76^ and the paediatric drug clearance data is less attainable, which is probably due to the difficulties associated with conducting paediatric clinical trials.^79, 80^ There exist model-based approaches for paediatric dose projection, such as weight-based dose prediction,^78^ Salisbury rule,^81^ paediatric dose prediction based on predicted clearance,^78^ and PBPK models^82^ as well as for predicting the total and renal clearance of renally secreted drugs in neonates.^83^ In future, these existing modelling techniques could be combined with the infant-WBMs allowing a detailed analysis of drug metabolism within the whole infant body potentially enabling a more accurate drug dosage determination and drug clearance prediction.^84^ To that end, PBPK models have already been developed to evaluate drug metabolism *in silico* for infants,^6^ which are aimed to complement the dose-extrapolation often performed for the determination of appropriate infant drug dosage.^7^ Genome-scale modelling, and WBMs, have already been connected with PBPK modelling, demonstrating the feasibility and value of such hybrid modelling approaches.^85–88^

## Conclusion

In summary, the presented resource of 360 infant-WBMs provides valuable insights into infant metabolism at an organ-resolved level and captures a holistic view of the metabolic processes occurring in infants, by considering the unique energy and dietary requirements as well as growth patterns specific to this population. They hold the promise for personalised medicine, as they could be a first step to a digital metabolic twin for newborn and infant metabolism for personalised systematic simulations and treatment planning.

## Methods

### 1 Generation of the infant-WBMs

The female and male infant-WBMs were developed using the female and male adult WBMs,^8^ respectively, as starting points. The adult WBMs are metabolic network reconstructions that have been assembled using organ-specific information from literature as well as omics data. These sex-specific reconstructions represent whole-body organ-resolved metabolism with over 80,000 biochemical reactions and capture the metabolism of 28 organs for the male and 30 organs for the female models. The adult WBMs have been parameterised using physiological, dietary, and metabolomic data.^8^ In the following, we delineate the changes that were made to the adult WBMs to derive the male and female infant-WBMs.

#### 1.1 Organ-specific biomass growth reactions

We assumed that all infant organs grow. Hence, organ-specific biomass growth reactions were formulated and added for each organ. A biomass growth reaction accounts for all known components required to replicate the cells of an organ.^13, 30^ These biomass growth reactions were modelled based on the biomass maintenance reactions that were already present in the adult WBMs, but now also included the molecular and energetic requirements for replication. Note that no changes were done for organs that contained already a biomass growth reaction (e.g., the skin). The addition of the organ-specific biomass growth reactions required the addition of further reactions to the WBMs to allow the biomass constituents to be transported into the correct compartment (Table S15). For instance, transport reactions for dNTPs from the cytosol to the nucleus needed to be added to allow for DNA replication requirements. A total of 236 and 259 reactions were added to the male and female infant-WBMs, respectively, resulting in 83,149 reactions in the male infant-WBMs and 85,662 in the female WBMs.

#### 1.2 Whole-body biomass reaction and organ weights

The whole-body biomass reaction represents the material and energy (ATP) required to maintain the non-metabolic cellular functions of the body and has been constructed based on the fractional weight contribution for each organ for a reference man or woman.^8^ Accordingly, the whole-body biomass reaction in the infant-WBMs was reformulated based on the age-dependent fractional organ weight distributions (Table S1), which were obtained from the literature.^6^ In the absence of infant-specific information, the relative weight of the female and male adult WBM^8^ was applied and scaled based on the infant’s body weight (Table S1). Note that for the female infant-WBMs, the organ weight of the uterus was further reduced, instead of using the relative value derived from the female adult model, for a more accurate representation (Table S1 (A)).

#### 1.3 Blood flow rates

Additionally, the organ blood flow rates were updated for newborns and infants (Table S1 (B), (C)) based on the infant blood flow rates^6^ and scaled by the infant’s organ weights (Table S1 (A)).^6^ These organ-specific blood flow rates and plasma metabolite concentration ranges, derived from metabolite concentrations obtained from the Human Metabolome Database (HMDB),^8, 89^ were used to define the lower bounds on the metabolite transport reactions from the blood compartment into the different organs for each reaction (see also equation 1, 2). For the kidney, we assumed that it filters 20% of blood plasma. We set the lower bound and upper bound for kidney filtration for each metabolite by multiplying the healthy plasma metabolite concentration ranges^89^ with the glomerular filtration rate^8^ (see also 2).

#### 1.4 Physiological parameters

We used phenomenological models specific for infants as a function of weight and age (Table S2) to obtain the following physiological parameters: the heart rate, stroke volume, cardiac output, hematocrit value, creatinine in urine, urine flow rate and glomerular filtration rate (Table S2). The cerebrospinal fluid (CSF) flow rate was approximated as

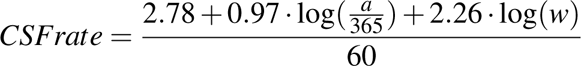

given in *ml/min* per day, where *a* is the age in days and *w* is the body weight in kg.^90^ In the original phenomenological model, a factor of 2.23 was subtracted for girls. However, for simplicity, we used this phenomenological model for both sex-specific infant-WBMs.

#### 1.5 Water balance

The urine excretion in infants has been approximated as

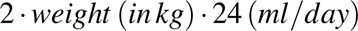

for their second day of life,^39^ which corresponds to an excretion of 158 ml in the urine of a 3.3 kg infant. This value corresponds to ≈10 % of the urine excretion of the male adult WBM, which has been estimated to be 1, 400 ml per day.^8^ Hence, we approximated the water loss through faeces, sweat, and air to be 10% of the adult WBMs on the second day of life. Based on this assumption, the amount of excreted water was increased daily depending on the daily water intake from the breast milk diet (Table S12 (A)). Additionally, we wanted to allow for a ± 10% margin on each day. Consequently, the upper bounds *ub_i_* and lower bounds *lb_i_* of the corresponding water exchange reactions *i* were based on the bound *b_i_* obtained from the adult WBMs^8^ and set as

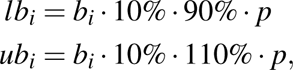

where *p* is the milk intake percentage (Table S12 (A)).

#### 1.6 Breast-milk based *in silico* diet composition during the first six months of life

We assumed the infant to be exclusively breastfed for the first six months of its life, which is in agreement with the recommendation by the WHO.^34^ The amount of daily milk intake was based on the milk model from the STIG-met publication^24^ and agrees with mean values of Swedish babies at 1, 2, 3, and 6 months of age.^91, 92^ We only increased the milk intake on the second day from 100 g to 225 g per day since on their second day of life newborns consume 22-27 ml milk per feed at 8-10 feeds per day;^93^ otherwise, the daily milk intake was not altered from the published milk model.^24^

The breast milk composition was derived from the human milk decomposition available at the Virtual Metabolic Human database (Table S9, https://www.vmh.life/).^35^ The obtained diet formulation consisted of approximately 87.5% water, which agrees with studies analysing the components of human breast milk.^94^ However, the original formulation did not contain lactose, which has been shown to account for approximately 7% of the breast milk.^95^ Hence, we added 7% lactose to the breast milk diet (Table S9). The original breast milk diet formulation was further modified to ensure the feasibility of the linear problem in FBA^26^ as well as adequate flux through the whole-body biomass reaction (see Table S9 and section). Moreover, to enable the growth of the infant-WBMs, the uptake bounds of dietary fluxes in the model of 12 metabolites had to be increased. These metabolites included six essential amino acids (L-methionine, L-isoleucine, L-valine, L-phenylalanine, L-threonine, L-leucine) and six other metabolites (choline, phosphatidylethanolamine, homocitrulline, D-glucose, thiamin monophosphate, guanidinoacetic acid) (Table S9). Furthermore, a low phosphate concentration in the diet had a growth-stunting influence on the model. We decreased the L-lysine concentration every month starting from two months, as it is an increasing factor for growth in the models (Table S5). This change is in agreement with L-lysine concentrations decreasing in breast milk after two weeks of lactation.^96^ Furthermore, for the female infant-WBMs, the dietary intake of L-cysteine was adapted as it presented another growth-limiting factor in the models (Table S5).

#### 1.7 Brain development

The brain development and its corresponding glucose demand are essential for the infant’s development and hence, for the infant-WBMs. The additional time and energy required for learning and brain development have been proposed to explain the slow and prolonged pre-adult life stage in humans compared to other species, even in comparison to other primate and great ape standards.^97, 98^ The brain glucose uptake per infant per day has been predicted to be 19.7 grams (converted to 100 mmol of glucose using the molecular weight of glucose 180,156 g/mol) by summing cerebral, cerebellar, and brainstem glucose uptake.^97^ Eukaryotes can produce 30-32 mol ATP from 1 mol of glucose depending on how the energy equivalents (in the form of NADH) from the cytosol get into the mitochondrium. We assumed that the glycerine-3-phosphate shuttle is used to produce 30 mol ATP from 1 g glucose.^99^ We integrated the energy requirement for the infant brain development into the infant-WBM by adapting the lower bound of the brain ATP demand reaction (VMH ID: Brain DM atp c). Specifically, the lower bound was set to be 100 mmol · 30 = 3, 000 mmol for the infant-WBMs at day one. This lower bound was increased every day, first steeply for two months and followed by a slower increase in accordance with the body growth of an infant (Figure 5 (blue line), Table S14).

**Figure 5:**
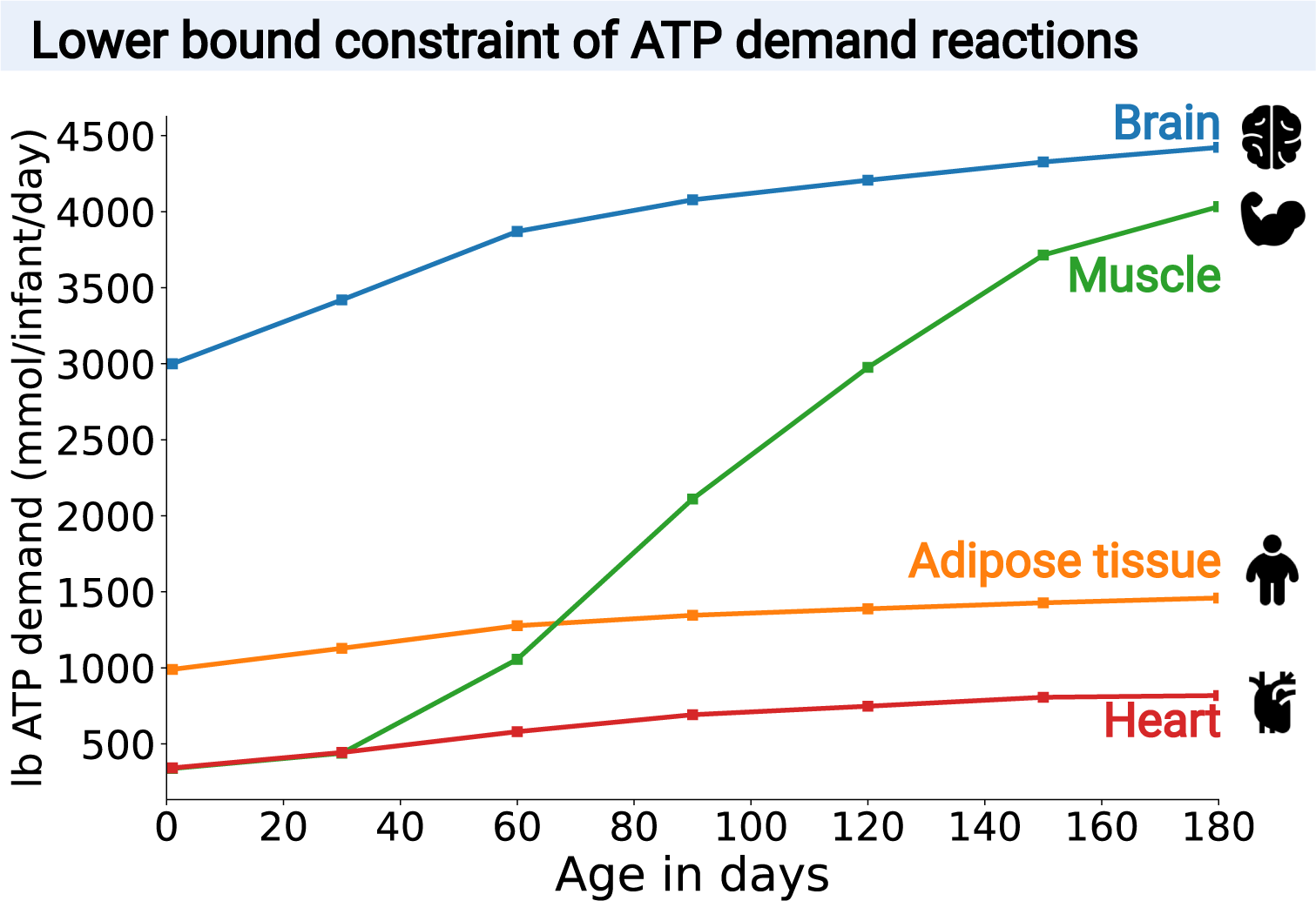
Energy balance. Lower bounds (lb) on ATP demand reactions in brain, heart, muscle and adipose tissue in mmol/day/person during the first six months of a male infant’s life.

#### 1.8 Thermoregulation

Thermoregulation of the human body refers to the ability to balance heat production and loss, ensuring the maintenance of the body temperature within a normal range. It holds particular significance for infants due to several factors: 1. infants possess less insulating fat compared to adults, 2. they have a relatively high surface-area-to-mass ratio, 3. a high head-to-body size ratio, and 4. an underdeveloped metabolic mechanism to respond effectively to thermal stress (e.g., no shivering).^100, 101^ Usually heat production by shivering is not functional in infants younger than six months since muscles are relatively immature to produce heat.^102^ Therefore, the mechanism of non-shivering thermogenesis was used as the primary mechanism for thermoregulation, which allows the generation of heat without relying on muscle contractions.^32, 33^ Non-shivering thermogenesis involves heat production that is not associated with muscular activity but is generated mainly by the brown adipose tissue and also to a smaller extent in skeletal muscle, brain, liver, and white adipose tissue.^103^ The brown adipose tissue contains an abundance of mitochondria and thermoregulation is enabled by an uncoupling protein, i.e., thermogenin or uncoupling protein (UCP-1).^32, 33^

The majority of the fat content in infants is comprised of brown fat.^104^ For infants, it has been estimated that between 44%^105^ and 55%^100^ of the total heat production comes from brain metabolism and heat production. It has also been reported that in the brain, approximately 1/3 of the glucose consumed by the brain is released as heat, while the remaining portion is utilised to generate ATP.^106, 107^ As we required the brain to have a non-metabolic energy demand of 3,000 mmol ATP per infant per day at age one day, we assumed that a minimum of a 1/3 of this value would correspond to the reported 1/3 of consumed glucose, even though the real value is likely higher as the overall metabolic and non-metabolic brain energy production exceeds 3,000 mmol ATP per infant per day in the models. Consequently, we set the lower bound of the adipose tissue energy demand reaction (VMH ID: Adipocytes DM atp c) to 3, 000 · 1*/*3 = 1, 000 mmol ATP per infant per day at age one day, and increased proportionally with the brain’s energy demand every day (Figure 5 (orange line), Table S14), to represent the near equal contribution of brain and brown fat to the infant’s heat production.

#### 1.9 Heart

In the adult WBMs, the heart ATP demand has been estimated to be 6,000 mmol ATP per day per adult with a heart weight of 330 g.^8^ In the absence of more specific data, we estimated the heart energy demand in the infant-WBMs based on the relative heart weight of the infant. As at birth, the heart weight is ≈20 grams,^6^ which corresponds to 6% of the adult heart weight, the lower bound on the heart ATP demand (VMH ID: Heart DM atp c) was set to 6, 000 · 6% = 360 mmol ATP on day one and increased daily in accordance with the growth of the heart weight (Figure 5 (red line), Table S14).

#### 1.10 Muscular activity

Another essential, energy-demanding mechanism in the infant’s body is the energy expenditure for physical activity. For this, we used a published activity model^24^ to estimate the physical activity. This activity model accounts for the difference between sleeping energy expenditure and total energy expenditure. The energy expenditure varies with age and has been determined to be 4.2 kcal/kg in surgically newborns,^108^ 10 kcal/kg in 3-month-old infants,^109^ and 14.4 kcal/kg in 4-6 month-old infants.^110^ To reflect this variation in the activity model the value of 14.4 kcal/kg is multiplied with a factor α ∈ [0, 1], which is based on estimated changes in physical activity from literature interpolated by a polynomial function of degree 2.^24, 111^ The energy expenditure was then converted to 28.12 mmol ATP hydrolysed per kcal, which has been estimated by simulating the production of ATP from glucose with the known energy content of 4 kcal/g.^24^ Consequently, the resulting value was multiplied by the infant’s body weight, leading to the calculated energy expenditure for physical activity of 328 - 3,515 mmol/day/infant ATP in the female models and 338 - 4,033 mmol/day/infant ATP in the male models, depending on age (Table S14). In the infant-WBMs, this calculated activity expenditure in ATP was used to set the lower bound of the muscle ATP demand reaction (VMH ID: Muscle DM atp c, Figure 5 (green line), Table S14).

### 2 Infant-WBMs for newborn screening

#### 2.1 Newborn screening data

The newborn screening data used in this study was obtained from the newborn screening laboratory at UKHD, Germany, which screens about 20% of the newborns in Germany (i.e., about 140,000 newborns per year).^60^ The UKHD data protection officer ensured that the newborn screening data were anonymised and that data extraction and evaluation were in accordance with the European general data protection regulation (GDPR).

The newborn screening data set comprised >2 million unremarkable newborn screening profiles from male (51%) and female (49%) newborns born between 2002 and 2021. Information for 48 metabolite concentrations and five additional variables, i.e., sex, birth weight, age at dried blood spot sample, age at sample arrival, and gestational age, were available for each newborn. To ensure high data quality, we removed missing values and uninterpretable entries from the data set. Furthermore, the following ranges were defined to exclude newborn screening profiles with implausible values and preterm newborns: Birth weight: 1000 − 6000 g; gestational age: 38 − 42 weeks, age at dried blood spot sampling: 36 − 72 hours, age at sample arrival: 0 − 20 days and metabolite concentrations: 0 − 50, 000µ mol/l. From the cleaned data set, consisting of 798,221 male newborns, we obtained a uniformly random sampled subset of 10,000 newborn screening profiles of male newborns to personalise the male infant-WBM utilising the concentration of 29 metabolites (Table S3), sex, and birth weight from each newborn. Note that only 29/48 metabolite concentrations were used as the remaining metabolites did not map onto the infant-WBMs. A Wilcoxon rank sum test confirmed that the birth weights of the subset (10,000 newborns) and the birth weights of the cleaned data set of male newborns (798,221 newborns) stem from the same distribution (p=0.27). This test was performed using the Python library scipy.^112^

#### 2.2 Integration of newborn screening data

Personalised infant-WBMs were generated using the concentration values of the 29 metabolites (17 amino acids and 12 acylcarnitines (Table S3)), sex, and birth weight of each newborn in the sampled subset. To set the bounds on the fluxes depending on the metabolite concentrations *m_conc,m_ ∈* R*_≥_*_0_ for each metabolite *m*, the metabolite concentration boundaries were calculated based on the concentration and a coefficient of variation *c_m_ ∈* R*_≥_*_0_ determined by the UKHD newborn screening laboratory, reflecting day-to-day variability within the tandem mass spectrometry.^113, 114^ The range of the fluxes was constrained with the coefficient *x_m_* = min(*c_m_,* 0.1). Hence, the minimum *m_min,m_* and maximum *m_max,m_* were calculated for every metabolite concentrations as:

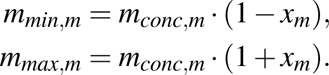

With *m_min,m_*and *m_max,m_*given in µmol/l blood. The lower bound *lb_m,kidney_* and upper bound *ub_m,kidney_*on the kidney metabolite uptake reaction fluxes were calculated using the glomerular filtration rate (GFR):

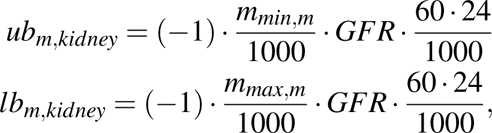

and are given in mmol/day/person. For all other organs, only the uptake of metabolites from the blood was constrained, and therefore the lower bounds on the uptake fluxes *lb_m,organ_* from the blood circulation into the individual organs was updated utilising the organ-specific plasma flow rate *PFR_organ_*,

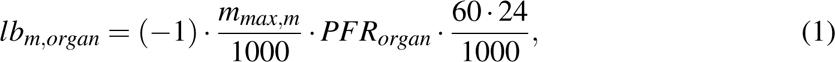

given in mmol/day/person. The organ-specific plasma flow rate was calculated as:

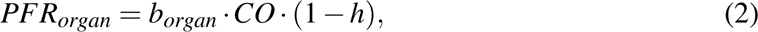

where *b_organ_* is the organ-, sex-, and age-specific blood flow percentage (Table S1 (C)), *CO* is the cardiac output and *h* is the hematocrit value. If no literature data was available for the blood flow percentage, it was set to the default value of 1%. The equations and flux relationships were established in Thiele et al.^8^

### 3 Simulation description

#### 3.1 Flux balance analysis

For the mathematical analysis and simulation of fluxes within a metabolic reconstruction, the reconstruction network is transformed into a stoichiometric matrix *S ∈* Z*^m×n^*, where the rows correspond to the *m* metabolites and the columns to the *n* reactions. The matrix entries *s_i j_* are assigned a stoichiometric coefficient if metabolite *i* takes part in reaction *j* and zero otherwise. The change of a metabolite concentration *x_k_* over time *t* is then represented by the *k*th row, while the computed flux value *v_i_* corresponds to *i*th entry in the flux vector *v* = (*v*_1_, …, *v_n_*)*^T^*. Resulting in the mass-balance equation for all metabolites as

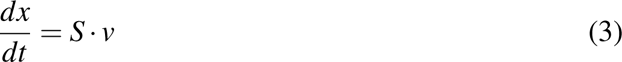

For the simulations, we use the COBRA approach,^12^ which assumes the simulated system to be at a steady state and, hence,

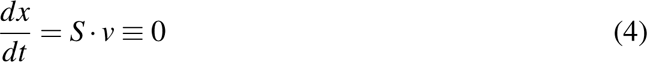

This assumption is then used as a constraint in FBA,^26^ which is a mathematical approach to minimise or maximise an objective function, e.g., biomass yield, through the metabolic network. The corresponding linear program (LP) is solved

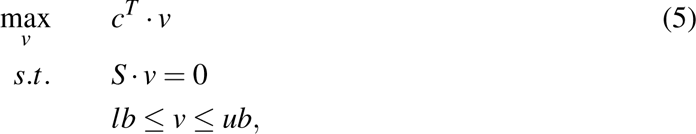

where *lb* and *ub* are the lower and upper bounds, respectively, on the flux vector *v* and they are based on specific data, such as physiochemical constraints, nutrient uptake rates, and enzyme reaction rates. The vector *c* is a vector of weights indicating which and how much each reaction *i* contributes to the objective function.^26^ By definition, the lower bound on an irreversible reaction was set to *lb* = 0 mmol/day/person and *ub >* 0 mmol/day/person.^8, 26^ For reversible reactions, negative flux through the reactions was allowed, *lb <* 0 mmol/day/person and the upper bound was set to *ub >* 0 mmol/day/person. The *lb* and *ub* of unconstrained reactions were set to the arbitrary values −1,000,000 mmol/day/person and 1,000,000 mmol/day/person, respectively. We defined uptake reactions, such as dietary intake, to have a negative flux value and excretion reactions, such as urine, faecal, air, and sweat excretion, to have a positive flux value.^8^

#### 3.2 Growth predictions

For the growth prediction, we iteratively applied FBA (*optimizeWBMmodel.m*) from the COBRA toolbox v3.0.^115^ Starting from day one, we maximised the flux through the whole-body biomass reaction (VMH ID: Whole body objective rxn) and used the computed flux solution to update the weight of the infant. For example, a computed flux value of 1.01 through the whole-body biomass reaction corresponds to a 1% gain in biomass, thus, increasing the body weight of the infant by 1%. Then, we used the new weight to create a model for the next day by updating all related constraints, and performed another FBA on this updated model, continuing like this until the age of six months.

As the matrix *S* has more rows than columns, the problem is under-determined, which leads to a polyhedral convex steady-state solution space containing all feasible steady-state solutions. However, to compare the fluxes through different reactions over time, we were interested in a unique flux solution to the problem. Therefore, we applied a second quadratic program (QP), where the Euclidean norm of the flux vector was minimised under the constraint that the optimal solution for the whole-body biomass reaction *v_whole body biomass_* was achieved:

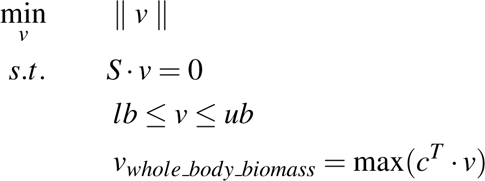

#### 3.3 Metabolic flux predictions within the infant body

We determined the aforementioned QP flux vector for each month (days 1, 30, 60, 90, 120, 150, and 180) in the infant’s life using the corresponding infant-WBMs. Then, we evaluated the flux predictions on these days for specific reaction fluxes of interest. For the water excretion, the flux values were retrieved and analysed for the diet, skin, urine, faeces, and air (VMH IDs: Diet EX h2o[d], Skin EX h2o[sw], EX h2o[u], EX h2o[fe], EX h2o[a]). For interpretability, we converted the flux through these reactions from mmol/person/day to ml/person/day by dividing the flux by 1000 and multiplying it with the molecular weight of water 18.02 g/mol since 1 g water = 1 ml water. For the evaluation of the ATP synthase, the flux values through the ATP synthase reaction of the brain (VMH ID: Brain ATPS4m), liver (VMH ID: Liver ATPS4m), muscle (VMH ID: Muscle ATPS4m), adipose tissue (VMH ID: Adipocytes ATPS4m), and heart (VMH ID: Heart ATPS4m) were compared for each month.

#### 3.4 Diet composition optimisation

The nutritional composition of the human milk diet varies over time and between mothers.^36, 67^ However, for comparability of our simulation results, and in the absence of personalised dietary data, we assumed the breast milk composition to be unchanged over time (except for the adaptations listed in the Method section 1.6). Some personalised infant-WBMs, which were personalised based on sex, birth weight, and newborn screening metabolomic data, were infeasible, i.e., no flux solution could be found that satisfied eq. 5. Similarly, some of the infant-WBMs evaluated at different points within the six-month time frame were also infeasible. For those infeasible models, we used a nutrition algorithm,^48^ implemented in the COBRA toolbox v3.0,^115^ to identify missing diet components that when added could render the LP problem feasible. In brief, the nutrition algorithm takes a given model and adds artificial reactions to the infant-WBMs, which mimic adding or removing dietary resources from the nutrition. These reactions produce artificial metabolites, called “Points”, which are also consumed by the biomass reaction, which is maximised. By doing so, the algorithm creates a connection between the biomass reaction and the diet components that will be either added or removed to/from the breast milk diet. Subsequently, the adapted infant-WBM is used in an FBA, which minimises the flux of points out of the model and a solution is determined such that dietary changes are minimised and the biomass reaction flux is maximised.^48^ Subsequently, the adapted infant-WBM is used in an FBA, which minimises the flux of points out of the model and a solution is determined such that dietary changes are minimised and the biomass reaction flux is maximised.^48^ We then evaluated the suggested dietary component additions against the literature and added compounds as needed (Table S5, Method section 1.6).

#### 3.5 Analysis of Inherited Metabolic Diseases

For further evaluation of the infant-WBMs, we predicted known biomarkers of five IMDs for three time points in the infant’s life and compared this to the prediction of the adult WBM.^8^ Usually, biomarkers are clinically measured either in the blood, urine, or CSF. Here, we predicted the flux through the biomarker in the blood compartment. We used the method established in Thiele et al^8^ encoded with the function *performIEMAnalysis.m* in the COBRA Toolbox v3.0.^115^ Briefly, for the simulation of any IMD, the reactions *k* = (*k*_1_, …, *k_n_*)*^T^* associated with the defect gene in any organ are identified. A dummy reaction *v_dummy_* is added to the infant-WBM, summing the equal contribution of each reaction *k_i_*. The flux through *v_dummy_* is then maximised. The maximal possible value *z* for *v_dummy_* is used to set the lower bound on *v_dummy_* to 75% of this value. Subsequently, for each biomarker metabolite in the blood compartment the demand reaction *v_demand_* is added and maximised. The corresponding LP problem is:

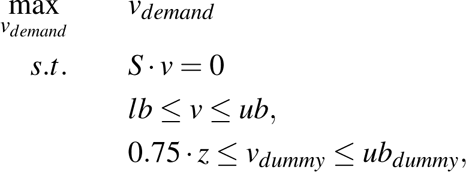

where *S* and *v* correspond to the stoichiometric matrix and flux vector, respectively, where the reactions *v_dummy_* and *v_demand_* are added to the original model. For the knock-out (disease) infant-WBM, the lower and upper bound of *v_dummy_* is set to 0, and the maximisation through each biomarker metabolite demand reaction *v_demand_* is determined.

Finally, the obtained fluxes within the biomarker reaction of the wild-type model *v_WT_* and the diseased model *v_D_* are compared. The relative flux increase is calculated as

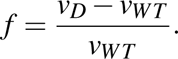

Using this equation, we could compare the relative predicted flux increase for any biomarker of interest for a specific IMD (Table S10 (B), (C)).

All simulations were carried out using Matlab 2020b (Mathworks, Inc.) as a programming environment, Ilog cplex as a linear and quadratic programming solver (IBM, Inc), and the COBRA Toolbox v3.0.^115^ All code will be part of the COBRA Toolbox v3.0 (https://github.com/opencobra/cobratoolbox/) and https://github.com/ThieleLab.

## Acknowledegments

The authors acknowledge the support of the Informatics for Life project funded by the Klaus Tschira Foundation, the financial support by the Dietmar Hopp Foundation, St. Leon Rot, Germany (grant numbers 2311221, 1DH2011117 and 1DH1911376), the financial support by the European Research Council (ERC) under the European Union’s Horizon 2020 research and innovation programme (#757922), the Science Foundation Ireland under Grant number 12/RC/2273-P2, and the Horizon Europe grant Recon4IMD (#101080997). The present work was also supported by the Helmholtz Association under the joint research school HIDSS4Health – Helmholtz Information and Data Science School for Health.

## Author contribution

Conceptualisation E.Z. and I.T.; Methodology E.Z. and I.T.; Software E.Z. and I.T.; Data Curation E.Z., U.M., J.G.O., and I.T.; Writing – Original Draft E.Z. and I.T.; Writing – Review & Editing E.Z., U.M., S.K., V.H., and I.T.; Supervision G.F.H., S.K., V.H., and I.T.; Funding Acquisition G.F.H., S.K., V.H., and I.T.

